# Modeling slow-processing of toxin messenger RNAs in type-I Toxin-Antitoxin systems: post-segregational killing and noise filtering

**DOI:** 10.1101/407288

**Authors:** Yusuke Himeoka, Namiko Mitarai

## Abstract

In type-I toxin-antitoxin (TA) systems, the action of growth-inhibiting toxin proteins is counteracted by the antitoxin small RNAs (sRNAs) that prevent the translation of toxin messenger RNAs (mRNAs). When a TA module is encoded on a plasmid, the short lifetime of antitoxin sRNA compared to toxin mRNAs mediates post-segregational killing (PSK) that contribute the plasmid maintenance, while some of the chromosomal encoded TA loci have been reported to contribute to persister formation in response to a specific upstream signal. Some of the well studied type-I TA systems such as *hok/sok* are known to have a rather complex regulatory mechanism. Transcribed full-length toxin mRNAs fold such that the ribosome binding site is not accessible and hence cannot be translated. The mRNAs are slowly processed by RNases, and the truncated mRNAs can be either translated or bound by antitoxin sRNA to be quickly degraded. We analyze the role of this extra processing by a mathematical model. We first consider the PSK scenario, and demonstrate that the extra processing compatibly ensures the high toxin expression upon complete plasmid loss, without inducing toxin expression upon acquisition of a plasmid or decrease of plasmid number to a non-zero number. We further show that the extra processing help filtering the transcription noise, avoiding random activation of toxins in transcriptionally regulated TA systems as seen in chromosomal ones. The present model highlights impacts of the slow processing reaction, offering insights on why the slow processing reactions are commonly identified in multiple type-I TA systems.

## 1. Introduction

Toxin-Antitoxin (TA) systems are ubiquitous among free-living prokaryotes [1]. Toxin proteins in the free form typically interfere with the cellular growth process, making the cells to be dormant or even killing the cell, while antitoxins neutralize the toxins [2–4]. The type-I TA module, where the antitoxin gene is transcribed to produce regulatory non-coding small RNAs (sRNAs) that prevent the translation of the toxin messenger RNAs (mRNAs), was first discovered as *hok/sok* of plasmid R1 in *Escherichia coli* [5]. The module contributes to the plasmid stabilization via post-segregational killing (PSK), causing the growth arrest or cell death for the cells that had lost the plasmid. Interestingly, *hok/sok* homologs, as well as other TA modules, have also been found as chromosomal genes [4]. The function of them is still under debate, but it has been found that *hokB/sokB* as well as *tisB/istR-1* on chromosome increase the persistence upon exposure to antibiotics by causing growth arrest to the subpopulation of the cells in response to specific upstream signaling [6,7].

There are several layers of molecular regulations that are important to achieve the functions mentioned above. Firstly, for PSK, the toxins should be activated when the plasmid is completely lost from the cell, but neither when the number of the plasmid decreased slightly with still keeping some in the cell, nor when the cell acquires the plasmid with a TA system for the first time. Secondly, for both PSK and signal-induced persistence, random activations of toxins at unwanted timing should be avoided.

One of the main mechanisms for the activation of the toxins upon plasmid loss has been attributed to the short lifetime of free antitoxin sRNAs compared to toxin mRNAs [8]. When the plasmid is lost, the antitoxin sRNAs disappear quicker than toxin mRNAs, allowing the toxins to be translated. Theoretically, this lifetime difference is possible in different forms. Recently, it has been shown by a mathematical model that, if the lifetime of the antitoxin sRNA-toxin mRNA duplex is long enough compared to the free sRNA and the complex can dissociate slowly, the toxins can be activated upon the plasmid loss [9]. However, for example in *hok/sok* of plasmid R1, the duplex is degraded rapidly [10, 11]. Instead, it has been found that, the full-length, stable toxin mRNAs form a secondary structure that hinders ribosome binding site. In order for the translation to occur, the 3’-end of the full-length mRNA needs to be properly processed. It is the translatable truncated mRNA that can be bound by antitoxin sRNA and degraded quickly (figure 1). The processing of the 3’-end is relatively slow, allowing some of the full-length mRNAs pooled in the cell. Upon plasmid loss, sRNAs disappear quickly, and the full-length mRNAs in the pool will be truncated and expressed, resulting in the toxin activation [12]. This slow processing of full-length toxin mRNAs into translatable truncated form appears to be a rather robust regulatory mechanism of a type-I TA system, since all the regulatory elements are found in *hokB/sokB* locus on *E. coli* chromosome [13,14], and also an almost completely equivalent mRNA processing was found in another type-I TA system, *AapA1/IsoA1* of *Helicobacter pylori* [15].

**Figure 1.**
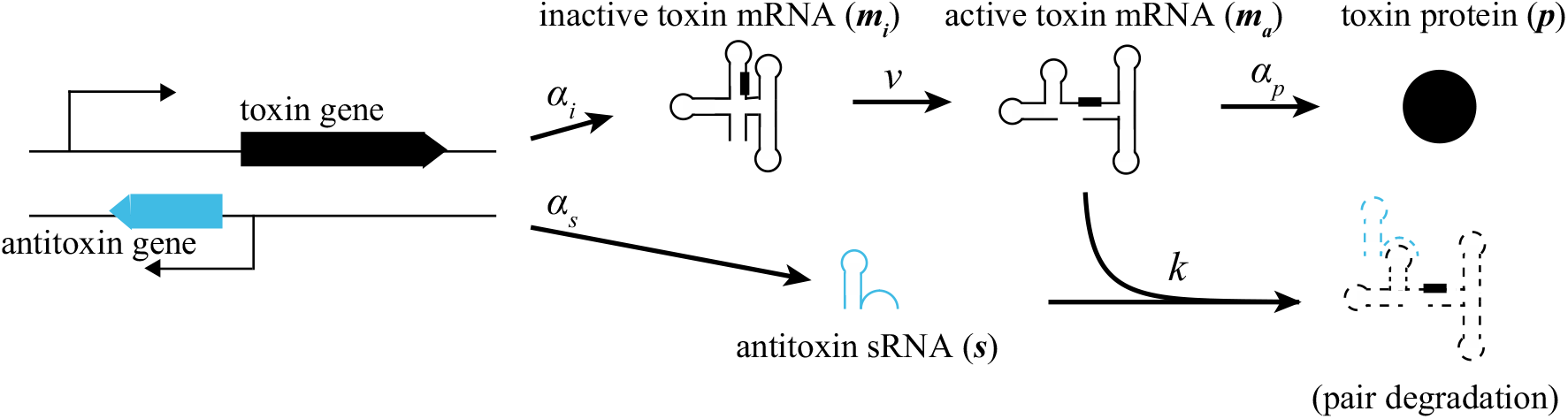
A schematic illustration of the model. The toxin mRNA and antitoxin sRNA are transcribed from the corresponding genes (at the rate α¾ and *α_s_*, respectively), whereas the toxin mRNA is not translatable because the ribosome binding site (represented by the black boxes on the mRNAs) is hindered by the secondary structure of mRNA (inactive toxin mRNA). The following processing reaction (at the rate *v*) changes the structure of toxin mRNA and the ribosomes can bind to it (active toxin mRNA). The toxin protein is translated from the active toxin mRNA (at the rate *α_p_*). The antitoxin sRNA interacts with the active toxin mRNA leading to the pair degradation (at the rate *k*). Each component is spontaneously degraded in at the rate *β*_*_, whereas the spontaneous degradations are not depicted in the illustration to avoid a complication of the figure.

The slow-processing of toxin mRNA is relatively complex, but the repeated appearance of the mechanism suggests that there are some benefits in this way of regulation. In this paper, we explore the possible advantage of the slow-processing mechanism in type-I TA system by using a mathematical model. We first consider PSK, and we show that the mechanism can suppress the toxin expression very tightly as long as the cell contains the gene, while upon the plasmid loss the mechanism can allow a rather high peak of the toxin expression. Furthermore, the system can keep the toxin expression low upon decrease of the plasmid number to non-zero number, and also avoid the expression of the toxins upon acquisition of the gene. We then analyze the fluctuation of the toxin expression when the gene copy number is a constant. We show that the slow processing contributes significantly in a reduction of noise, avoiding the unexpected activation of the toxin in unstressed cells.

## 2. Model

We construct a model for type-I toxin-antitoxin systems with the processing of full-length toxin mRNA (figure 1). Hereafter, we refer the full-length toxin mRNA and truncated toxin mRNA as “inactive toxin mRNA” and “active toxin mRNA” based on the translatability, respectively. Already known molecular mechanisms of the type-I TA systems [5, 11, 12, 16–18] leads to a simple model consists of the concentrations of the inactive toxin mRNA (*m_i_*), active toxin mRNA (*m_a_*), antitoxin sRNA (*s*), and toxin protein (*p*) given by

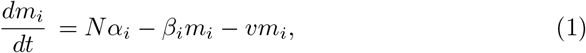

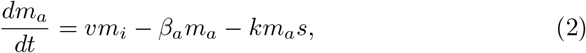

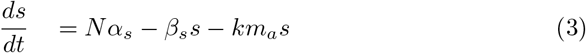

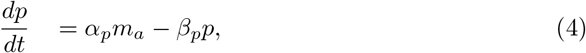

where *N*, *α_i_*, *α_s_*, *α_p_*, *β_i_*, *β_a_*, *β_s_*, and *β_p_* represents the number of plasmids, synthesis rate of inactive toxin mRNA, antitoxin sRNA, toxin protein, the spontaneous degradation or dilution rate of inactive toxin mRNA, active toxin mRNA, antitoxin sRNA, and toxin protein, respectively. Transcribed inactive toxin mRNAs are not translatable and do not interact with antitoxin sRNA, while they are slowly processed into the active toxin mRNA at rate v. Active toxin mRNAs (*m_a_*) are translationally active and interact with antitoxin sRNA. An active toxin mRNA and an antitoxin sRNA forms a RNA duplex. Because it is reported that the RNA duplex is quickly degraded [10, 11], we omit the dissociation reaction, as has been done often in the modeling of sRNA regulations [19–22]. Active free toxin mRNAs are translated, and toxin proteins are synthesized at rate *α_p_* · *m_a_*. All chemical components spontaneously disappear at rate β_*_. Here, we incorporate the effect of dilution due to the volume growth of the cell into the spontaneous degradation rate.

In the result sections, we use the following parameters values as a default. We consider *E. coli* of volume 1*μm*^3^ as a typical cell, therefore 1 nM of molecule concentration corresponds to one molecule per cell [23]. The most of default parameter values are determined based on the studies for *hok/sok* system, which is the best characterized type-I TA system. The hok mRNA is stable, while the quick degradation (halflife ≈ 30 seconds) is reported for sok sRNA [16,17]. Additionally, the slow processing reaction of the full-length *hok* mRNA into truncated *hok* mRNA was around an hour [11]. Based on these reports, we set *β_i_* = *β_a_* = ln2/30 (min^−1^) (the effect of dilution with the doubling time 30min), *β_s_* = ln2/0.5 (min^−1^), and *v* = ln 2/60(min^−1^). For the toxin proteins, we set parameter values which are typically used for the study of the protein dynamics [24]. The protein synthesis rate per 1nM of mRNA is set to be *α_p_* = 5.0(min^−1^), and we set *β_p_* = ln 2/30(min^−1^), as an effect of the dilution. We set the second-order duplex formation rate to be *k* = 6.0 (nM·min^−1^) based on the theoretically inferred parameter range [9]. The results shown below are not altered qualitatively as long as *k* ≥ 0.1 nM·min^−1^ holds ‡.

The production rates of two RNAs have not yet been measured. Thus, we use values for other mRNA/sRNA systems reported in [19], where the transcription rates are inferred as in the order of 10^−3^ ~ 1 (nM · min^−1^) for mRNAs and 10^−1^ ~ 10 (nM · min^−1^) for sRNAs. We use α_s_ = 10.0 (nM · min^−1^), α_i_ = 0.5 (nM · min^−1^) as a default parameter values, whereas we also simulate large parameter range to show the robustness of the results.

## 3. Results

### 3.1. Accumulation of toxin mRNAs without toxin expression in the steady state

For the PSK mechanism to work, the steady state with *N* > 0 should ensure very low toxin protein level, and at the same time enough level of toxin mRNAs should exist so that the toxin can be expressed without additional transcription upon plasmid loss. In the present model, the steady state concentration of the full length toxin mRNAs is given by

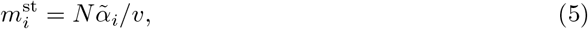

with

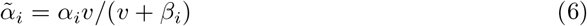

representing the effective production rate of the active toxin mRNA in the steady state. The steady state level of other quantities can also be exactly calculated (Appendix A), but here we present a simpler approximated solution. Since the duplex formation is the dominant part of the degradation of active toxin mRNA under the existence of the plasmid copies, we can ignore the spontaneous degradation / dilution of the active mRNA *m_a_* in (2). Under this approximation with the condition that antitoxin sRNA production rate is higher than that of the active mRNA 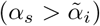, we get

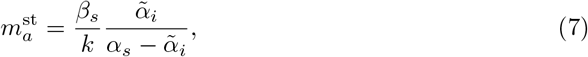

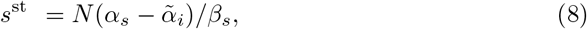

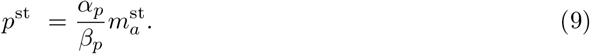

Note that the steady state concentrations of active toxin mRNA and toxin protein are independent of the plasmid copy number *N* because the increase of production rate of active toxin mRNA is completely compensated by the increased production of antitoxin sRNA under this approximation. The antitoxin sRNA concentration *s*^st^ is simply proportional to the difference between the two production rates, 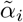 and *α_s_*. Since the inactive toxin mRNA does not interact with sRNAs, it is possible to keep 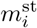 high and keep active toxin mRNA concentration 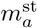 and hence the toxin protein concentration *p*^st^ low, by having large enough *α_s_*.

### 3.2. The slow processing reaction helps to satisfy the conflicting demand before and, after plasmid loss

Figure 2a shows temporal responses of the concentration of the toxin protein for the acquisition (*N* = 0 → 1) and loss (*N* =1 → 0) of a plasmid. At *t* = −300 minutes, the cell acquires the plasmid and starts transcribing the toxin mRNA and antitoxin sRNA coded on the plasmid. Then, the toxin protein is produced subsequently and its concentration reaches the steady values. The increase of the toxin is slow and kept well below 1 nM, because the slow processing of the inactive toxin mRNA gives enough time for the antitoxin sRNAs to be produced and prevents the translation of active mRNAs when they slowly appear.

**Figure 2.**
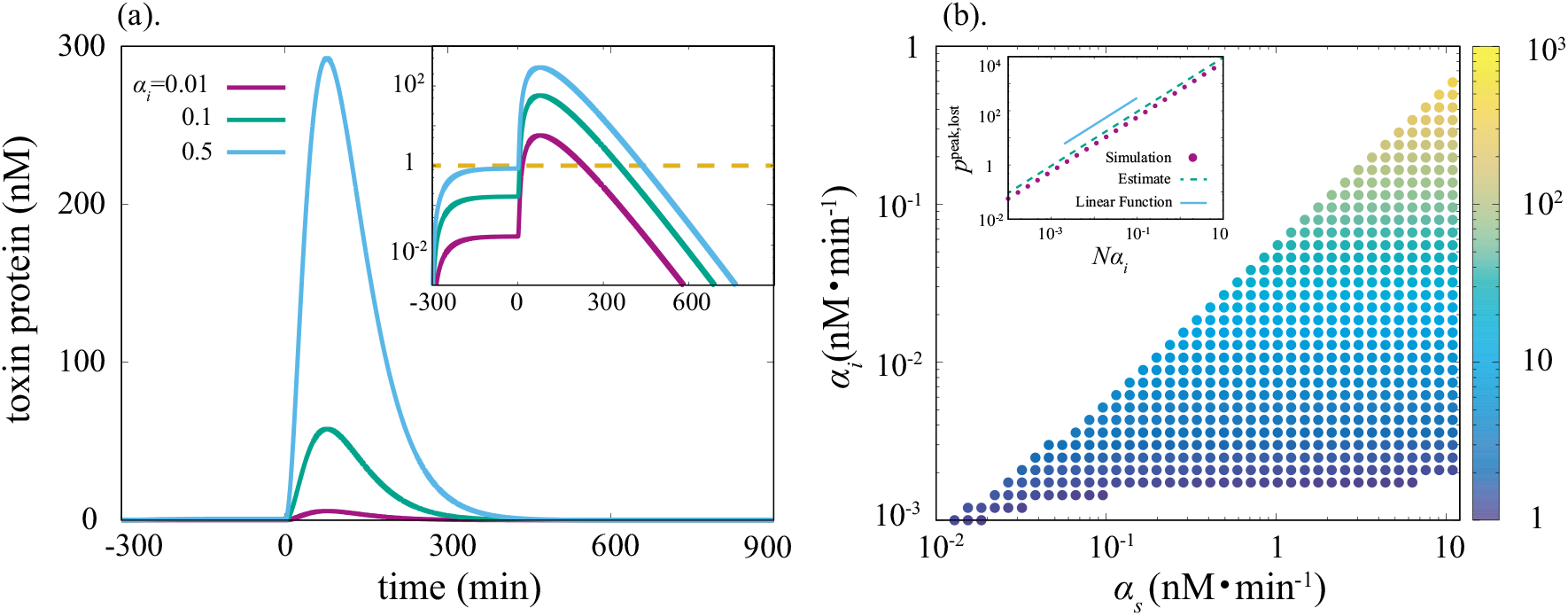
(a). Transient responses of the toxin protein concentrations to the plasmid acquisition(*t* = −300) and disappearance (*t* = 0) events. Inset: A semi-log representation of the concentration of toxin protein. (b). The parameter region of the production rate of the toxin mRNA and antitoxin sRNA in which the system works as PSK mechanism. Color indicates the concentration of the toxin protein at the peak after the plasmid loss (*N* = 1 → 0). Inset: the comparison between the estimated value (B.5) in Appendix B) and the numerical solution. The linear function is also plotted as a reference.

The plasmid loss occurs at *t* = 0. The transient response also comes from the delayed dynamics of the toxin mRNA. At the steady state with *N* > 0, the production rate of the active toxin mRNA is much smaller than the transcription rate of antitoxin sRNA. When the transcription of these two RNAs stops due to the plasmid loss, the antitoxin sRNAs are quickly degraded. On the other hand, the active toxin mRNA is still produced by the processing of the inactive toxin mRNA. This difference between the kinetics of the toxin mRNA and the antitoxin sRNA implements the drastic increase of the toxin protein after the plasmid loss. In other words, after the plasmid loss, the inactive toxin mRNA and slow processing process work as a ‘reservoir” of active toxin mRNA.

We performed a parameter scanning to obtain a parameter set of *α_i_* and *α_s_* at which the steady concentration of the toxin protein is less than 1 nM (one molecule per cell) with one plasmid, whereas it exceeds 1nM after the plasmid loss. As shown in figure 2b, the present model can attain this condition in a wide range of the parameter space.

For PSK to work properly, the number of toxin protein should be lower than the certain threshold as long as the cell carries plasmid copies, while it should go beyond the threshold once the cell loses all plasmid copies. Although the threshold value is unknown, the value should be significantly higher than the steady concentration with plasmid and significantly lower than the peak concentration upon plasmid loss.

The slow processing reaction plays a crucial role to resolve these contradictory demands. For the quantitative characterization of the necessity of the slow processing reaction, we computed both the steady state and the peak concentration of the toxin protein as a function of the processing speed *v* (figure 3a). For small enough *v*, the peak value of the toxin protein concentration increases with *v*, and there is a large difference between the steady and the peak concentrations. As *v* increases further to approach the antitoxin sRNA degradation rate *β_s_* (set to order one), however, the peak concentration decreases and finally becomes equal to the steady-state concentration (i.e, no response to the plasmid loss event). This is because the inactive toxin mRNA no longer works as a “reservoir” of active toxin mRNA upon plasmid loss at large *v*. The steady concentration of the inactive toxin mRNA is given by 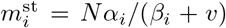 representing the capacity of the “toxin mRNA reservoir”, and it decreases with an increase of *v*. Note that the value of *v* also affects the parameter region of *α_i_* and *α_s_* in which PSK mechanism works (figure 3b).

**Figure 3.**
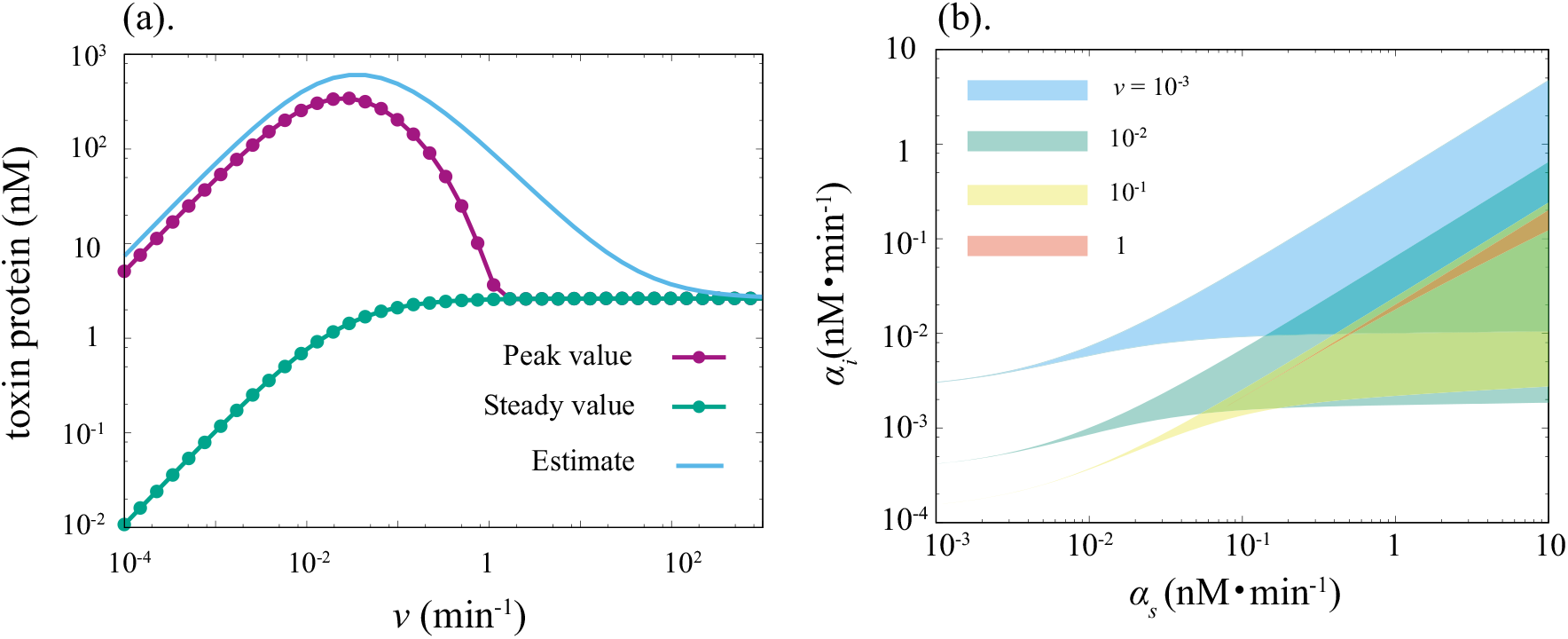
(a) The steady- and the peak concentration of the toxin protein is computed for several values of *v*. Up to *v* ~ 10^−2^/min, both the concentrations increase as *v* increase, while two values collapse at *v* ~ 1/min. (b) The parameter regions of the production rate of the toxin mRNA and antitoxin sRNA in which the system works as PSK mechanism.

The relationship between the capacity of the reservoir and the necessity of the slowness of the processing reaction can be analytically explored. In Appendix B, we analytically estimate the concentration of the active toxin mRNA at the maximum value during the transient response assuming *v* ≪ *β_s_*. This assumption gives us the estimate for the ratio between the peak concentration 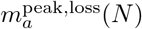 and the steady state concentration of the active toxin mRNA 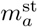 as

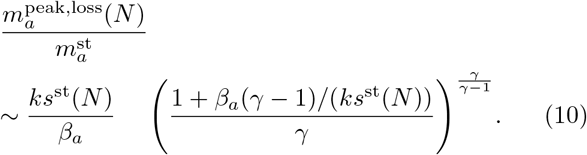

with a rescaled parameter *γ* = (*v* + *β_i_*)/*β_a_*. The estimated value is an increasing function of the intensity of the RNA duplex formation, *ks*^st^(*N*), as long as *γ* > 1 and *ks*^st^(*N*) > *β_a_* hold (Note that the latter condition is assumed to obtain the approximated solutions).

The peak concentration of the toxin protein is roughly estimated by multiplying 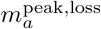 by *α_p_*/*β_p_*, and this is plotted in figure 3a against *v*. Although the time-scale separation between the active toxin mRNA and toxin protein is not clearly held with our default parameter set, the estimated value captures the characteristics of the concentration of the toxin protein reasonably well. Also, the estimated value can be simplified further by assuming *β_a_*(*γ* − 1)/(*ks*^st^(*N*)) ≪ 1 holds§. By omitting this term and substituting the approximated solutions (7) and (8) into (10), we obtain

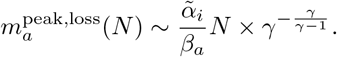

This simplified estimate shows that the peak concentration of the active toxin mRNA, and accordingly that of the toxin protein, linearly increase with the number of the lost plasmid copies. This linearity is numerically verified as shown in the inset of figure 2b.

### 3.3. Response to the decrease of plasmid number

For the plasmid maintenance, plasmid-free cells should be removed from the bacterial population, whereas cells should be protected from the host-killing mechanism as long as they carry the plasmid. In other words, a strong increase of the host-killing proteins should happen only when the plasmid copies are completely lost, but not when the number decreases to a non-zero number at the cell division.

Figure 4 shows the dynamics of the concentration of the toxin protein during sequential loss events of plasmids. We set the initial condition at *t* = 0 so that a cell is in the steady state with *N* = 10, and the cell loses *N*_dec_ plasmid(s) (*N*_dec_ = 1, 2, 5, and 10) simultaneously for every *τ* = 300 × *N*_dec_ minutes intervals. The cells lose all plasmids at the time *t* = 3000 minutes. As shown in figure 4(a) and (b), the peak values of the toxin protein at the every plasmid decrease events occurring *t* < 3000 minutes are at most 1nM level, while the response to the final event is in *μM* order. The peak values of the toxin protein concentration during the response to the plasmid decrease events are determined not by the absolute numbers of plasmid copies before and after the events, but by the ratio of the plasmid number between before and after the events. The peak value during the plasmid decrease event in which the number of plasmid copies changes *N* to *N*′ is analytically estimated by assuming *β_a_* = 0 as

**Figure 4.**
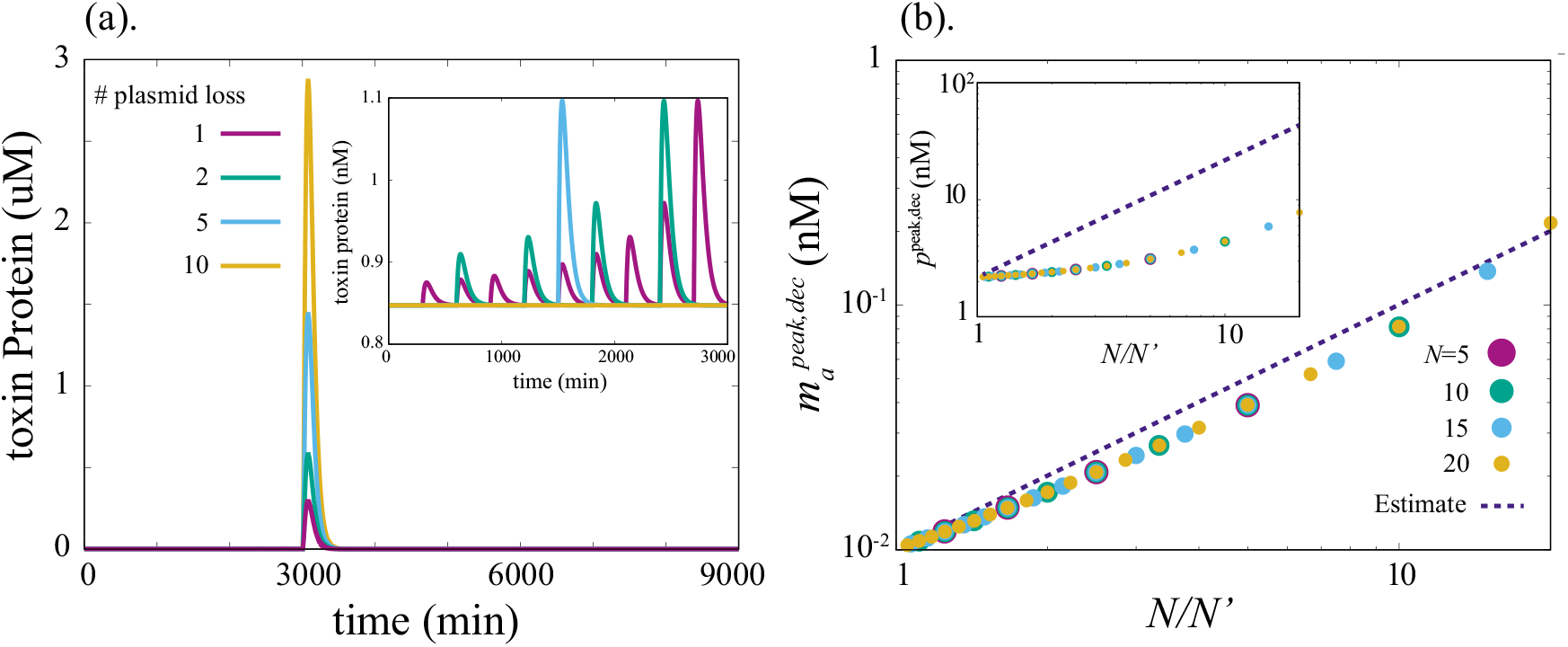
(a). Responses of the concentration of toxin protein to the decrease of plasmid copies. The decrease events of plasmid number occur at every fixed interval. The initial number of plasmid copies is set to be 10. The number of plasmid decrease for every event is indicated in legend, the interval of the decrease event for each *N*_dec_ value is set so that the cell loses all plasmids at *t* = 3000 (i.e the interval is 300(min) for purple line, and 3000(min) for orange line. Inset: The enlarged figure of the time series up to *t* = 3000. (b). A comparison between the estimated value (11) and numerical results of the peak concentration of the active toxin mRNA. Inset: A comparison between the estimated value and numerical results of the peak concentration of the toxin protein. Dots are obtained by computing the peak concentration of the active toxin mRNA (the toxin protein for inset) during the transient response of the plasmid decrease event (*N* → *N*′), and plotted against *N*/*N*′. The peak value is computed for all *N*′ values while less than *N* and larger than 0 for each color.)

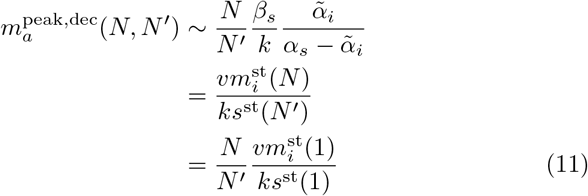

(Appendix D for detail), which is a reasonable estimate compared to the numerical results (figure 4b). Interestingly, the peak value depends not on the absolute number of the plasmid copies, but the fold change of the copy number, as shown in figure 4b ∥. The peak toxin protein concentration also depends on the fold change *N/N*′, but the actual value deviates from the estimate obtained by multiplying *α_p_/β_p_* to Eq. (11) (figure 4b inset) because the lack of the clear timescale separation between the active toxin mRNA and toxin protein affects the protein dynamics.

Clearly, our analytical estimates (10) and (11) show that an increase of the steady concentration of the antitoxin sRNA or the rate of the duplex formation (*k*) have only positive effect to the PSK mechanism: Enhancing the duplex formation not only magnifies the difference between the steady- and the peak concentration of the toxin protein in the loss of plasmid copies, but also makes the mechanism accurate so that the mechanism responds only if all plasmid copies are lost.

### 3.4. Slow processing as a noise reducer

Type-I TA systems are found not only in plasmid copies but also in bacterial chromosomes. The slow processing of inactive toxin mRNA into active mRNA is observed also for several chromosomal type-I TA systems such as *tisB/istR-1, dinQ/agrA*(*agrB*), and *AapA1/IsoA1* [4, 12, 15, 18, 25]. In this section, we study the role of the processing reaction for chromosomal TA systems.

The regulatory design of TA genes in the chromosomes is somewhat different from that of plasmid-borne TA systems. The toxin gene is repressed by a specific transcription factor under nonstressed conditions, whereas the repression level is down-regulated by a specific signal (for example, the expression of *tisB* gene is repressed by LexA protein, and the SOS signal facilitates the degradation of LexA protein to enhance the *tisB* expression [25–27]). Then, the toxin mRNA, and accordingly, the toxin protein is synthesized to interfere with the vital cellular processes and leads to the persister formation.

Recall that the slow processing of inactive toxin mRNA leads to the formation of the “reservoir”. Since large reservoirs of molecules typically contribute to reducing noise in stochastic chemical reaction systems, here we focused on the possible scenario that the processing reaction act as a noise reducer of the toxin protein, and performed stochastic simulations.

We constructed a model for chromosomal type-I TA systems with transcriptional regulation by slightly modifying our model for the plasmid-borne type-I TA systems: The number of plasmid copies N is considered as the number of TA genes on the chromosome, and it is set to unity. Also, the dynamics of the repression by repressor is modeled by introducing two distinct states (ON and OFF) into the toxin gene locus. In ON state, the toxin gene is not inhibited by the repressor, and the inactive toxin mRNA is produced at the rate *α_i_*, whereas there is no production of the inactive toxin mRNA in OFF state. The toxin gene transits ON and OFF states in the rates *l*_−_ (ON to OFF; typically increase with the repressor concentration) and *l*_+_ (OFF to ON; typically decrease with the binding strength of the repressor to the operator site); The average repression factor for the gene is then given by *l*_−_/(*l*_+_ + *l*_−_). For the quantitative characterization of the role of the processing reaction, a TA model which lacks the processing reaction is constructed additionally. The inactive toxin mRNA is eliminated from the model, and thus, the translatable toxin mRNA was transcribed from the toxin gene directly at the rate 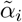 in (6) so that the translatable toxin mRNAs are produced at the same rate in the steady state for both of the models.

We compare the Fano factor (the ratio of the variance to the mean) of the number of toxin protein for with- and without processing. The Fano factor for each model is computed from the model via the Gillespie algorithm [28] (the detailed model equations are shown in Appendix E.). As shown in figure 5, the Fano factor of the model with the processing reaction is always smaller than that of without the processing reaction.

The transcription noise coming from the ON-OFF dynamics directly affects the probability of the toxin protein production in the model without slow-processing reaction. Therefore, if the number of the antitoxin sRNA is few by chance, the transition from OFF state to ON state can lead to the considerable amounts of the toxin protein production. In contrast, if the cell has the toxin mRNA is produced in inactive form and activation is slow, the fluctuation of the transcription of the toxin mRNA is buffered by a large number of the inactive toxin mRNA, and is averaged out over time by the slow processing reaction. In other words, the production rate of the active toxin mRNA is roughly approximated by *v* < *m_i_* >_1/*v*_ with < *m_i_* >_1/*v*_ being the time average of the inactive toxin mRNA for 1/*v* (min.). < *m_i_* >_1/*v*_ approaches a constant value as 1/*v* becomes longer, and the active toxin mRNA production approaches to a Poisson process, buffering the bursty production of inactive toxin mRNA.¶

The only difference between the models with and without slow processing is the existence of this ‘‘buffer” variable, and thus, the difference in the noise should be reflected in the production of the active toxin mRNA. Therefore, we analytically calculated the Fano factors *F_with_* and *F*_w/o_ of the number of the active toxin mRNA for simplified version of each model where the anitoxin sRNA is set to zero (For the model equation see Appendix F). The ratio of Fano factors is given by

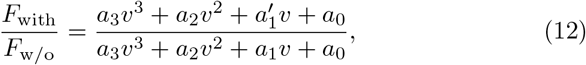

where *α_i_*’s and 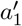 are constants independent of *v*. The detailed expression of these constants given in Appendix F show that all of these constants are positive and 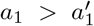 holds regardless of parameter values. This means 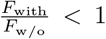, i.e., the combination of the inactive toxin mRNA and the slow processing reaction always reduces the fluctuations in the number of the active toxin mRNA. The expression (12) is plotted in figure 5b, demonstrating that it captures the noise reduction effect in the toxin protein in the full model.

**Figure 5.**
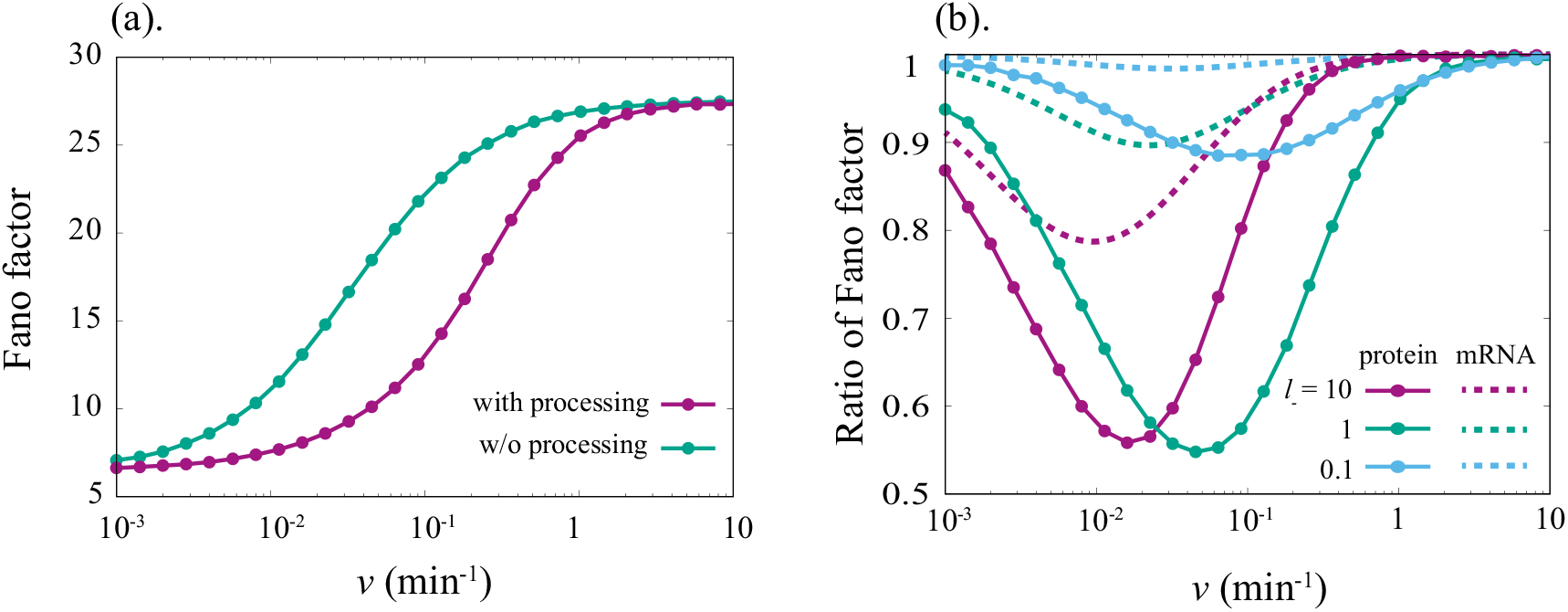
Comparisons of the Fano factor of the number of toxin protein between with- and without the processing reaction of the inactive toxin mRNA. (a). The Fano factors for the two models are plotted against the processing rate *v*. *l*_−_ is set to unity. (b). The ratio of the Fano factor with the processing reaction to that without processing reaction for several values of ON→OFF transition rate. The ratio of the Fano factor of the number of the active toxin mRNA without the effect of antitoxin sRNA (12) is also plotted against *v* (dashed lines). The parameter values are set to *α_i_* = *α_s_* = 1.0 and *l*_+_ = 10^−2^ (*l*_−_ values are shown in the plot) while others are same with the default values.

## 4. Discussion

We have constructed a simple model for the type-I TA system where the slow processing of the inactive toxin mRNA to a translatable form is explicitly taken into account. The analysis of the model highlighted an outstanding role of the untranslatable form of toxin mRNAs and slow processing reaction. Thanks to the time delay between the production of active toxin mRNA and that of antitoxin sRNA, the model attained contradicting demands needed for the PSK to work: the toxin protein is kept low concentration at the acquisition of plasmid and also as long as the cell carries plasmid copies, whereas a drastic increase of the toxin protein was achieved when the cell becomes plasmid-free. In addition, we have shown that such a strong increase of the toxin protein concentration does not take place when the number of plasmid copies decreases but the cell still carries at least one plasmid.

Having the inactive mRNA and slow processing reaction is not only beneficial for the plasmid-borne TA systems, but also for the TA systems on bacterial chromosomes. We introduced an ON/OFF kinetics of the toxin gene transcription into our model to emulate the noise from regulation of type-I TA systems on bacterial chromosomes. To highlight the role of the slow-processing, we compared the result with a simpler model where the transcribed mRNAs are already translatable and interact with antitoxin sRNAs. Stochastic simulations of the two models showed that the slow processing reaction reduces the stochastic burst of the toxin protein copy number.

In general, TA systems accomplish the need to produce toxins after the loss of the TA genes by the time-scale difference between the antitoxin life-time and the toxin life-time. The examples include not only type-I but also type-II TA systems, where the transient toxin excitation is accomplished by using the dissociable toxin-protein and antitoxin-protein complex as a reservoir of the active toxin proteins [4, 29, 30]. The uniqueness of the present model TA system is that the *inactive* toxin mRNA works as a non-toxic ‘reservoir”. If the translatable mRNAs should work as a non-toxic reservoir as a dissociable complex with the antitoxin sRNAs, the demand for high-peak of toxin at the plasmid loss requires relatively high concentration of the complex in a cell with the TA gene and the dissociation rate faster than the disappearance rate of the duplex [9]. Such requirements increase the chance of the toxin protein production before the plasmid loss. The situation is similar in the type-II systems where the toxic proteins need to be preserved in the reservoir before plasmid loss. The slow-processing mechanism neatly decouples the trade-off between the low chance of the toxin protein production in a cell with the TA genes and the high concentration of toxin level after the plasmid loss. The high level of the inactive mRNAs can be buffered easily by enough level of antitoxin sRNA concentration before plasmid loss, while after the plasmid loss the free sRNAs are degraded quickly to allow production of active toxin mRNAs. It should be noted, however, that in some of type-I TA systems the intermediate state has not been found [31,32]. Such systems may make use of the dissociable complex as a way to keep toxin mRNAs after plasmid loss [9].

The slow-processing of toxin mRNAs is commonly identified in a large variety of type-I TA systems even among those found in different classes of proteobacteria [15, 18, 25]. Thus far, the regulations by small noncoding RNAs have been extensively studied [9, 19−22], while the role of the intermediate state has not been highlighted theoretically. The present model provides an essential feature of the state offering insight to explore the factors which make the intermediate state a well-conserved feature.

## Acknowledgments

The authors thank Kenn Gerdes for fruitful discussions. This work was funded by the Danish National Research Foundation (BASP: DNRF120).

## Appendix A. steady solution

The exact steady solutions of the chemical species are straightforwardly calculated from (3), and given by

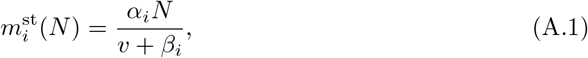

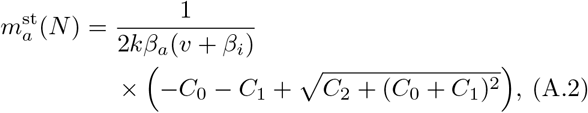

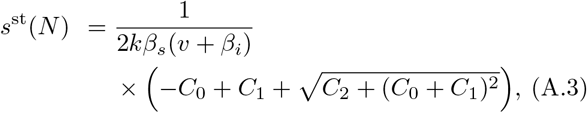

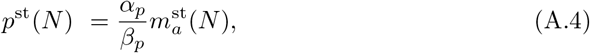

where

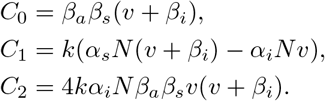

Note that the exact solution and the approximated solution for the inactive toxin mRNA are identical. By setting *β_a_* = 0, we get the approximated steady-state solutions presented in the main text.

## Appendix B. Analytical approximation of the peak concentration for a plasmid loss event

Assume that up to time *t* = 0, the cell has *N* copies plasmid, and the concentration of the molecules are the steady state values. Then, the copy number of the plasmid immediately drops to 0. We evaluate the approximate solution of the dynamics of the molecular species after *t* = 0. Here, we use the approximated steady solutions (7)–(9) obtained by assuming *β_a_* = 0 for calculations, whereas *β_a_* is not ignored for *t* > 0 because this approximation is no longer valid after the cell becomes plasmid-free.

First, we start with the dynamics of inactive toxin mRNA concentration. By knowing (A.1), the exact dynamics is given by

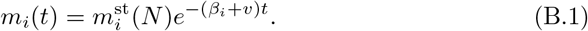

Note that the production of the antitoxin sRNA is halted for *t* > 0, and the spontaneous degradation has greater effect than the duplex formation to the decrease of antitoxin RNA concentration at the beginning of the relaxation because the concentration of active toxin mRNA is strongly suppressed under the existence of plasmid (*m_a_* ~ 0 at *t* = 0), and the timescale of the accumulation of the active toxin mRNA after the plasmid loss (~ 1/*υ*) is much longer than that of spontaneous degradation of antitoxin RNA (~ 1/*β_s_*). Thus, we can approximate the dynamics of the antitoxin sRNA concentration as

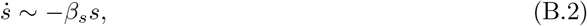

and it leads to *s*(*t*) ~ *s*^st^(*N*)*e*^−*β_s_t*^. The active toxin mRNA is rate-limiting substrate of the toxin:antitoxin duplex formation up to *t* ~ 1/*β_s_*. Thus, we can estimate the amount of active toxin mRNA which form the duplex with antitoxin RNA and degraded by assuming that the all toxin mRNA produced from the inactive toxin mRNA within the timescale forms duplex and degraded. The amount is calculated as

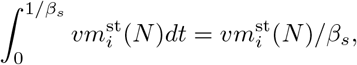

and the fraction of this value to the total production of active toxin mRNA during the relaxation timecourse is (*β_i_* + *υ*)/*β_s_*. With our default parameter settings, this fraction is about 2.5%. Therefore, the duplex formation has only minor effect to the peak value of the toxin protein, and thus, we assume that s reaches to zero instantaneously at *t* = 0.

Then, we get an approximated differential equation for *m_a_* as

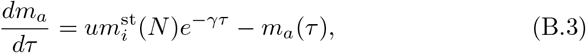

where we rescaled parameters such that *τ* = *β_a_t, u* = *υ/β_a_*, and *γ* = (*υ* + *β_i_*)/*β_a_*. The time-dependent solution of *m_a_* = *m_a_*(*τ*) can be obtained from this equation. By solving *dm_a_/dτ* = 0 for finite *τ*, the peak value of *m_a_* for the plasmid lost (*N* → 0) event is estimated as

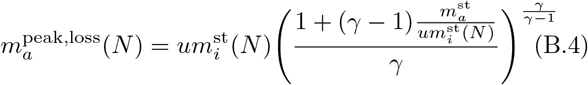

By dividing this expression by 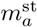 and using the relation 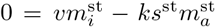, we get (10). The peak value of the toxin protein concentration is estimated as

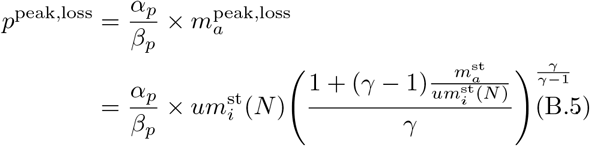

subsequently with the assumption of timescale separation between *p* and *m_a_*.

## Appendix C. Necessity of the processing reaction

To show the necessity of the processing reaction to generate a large gap between the concentration of protein at the steady-state and at the peak after the plasmid loss, we study a model derived by eliminating the processing reaction from the original model (3) in which the active toxin mRNA is produced at the rate 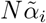 directly. Here we assume the rate of duplex formation is infinitely fast so that the results are obtained analytically. First, the steady-state concentrations of chemicals with *N* ≥ 1 plasmid copies (written as 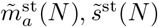, and 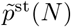) are given by

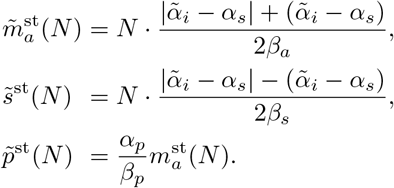

From the steady solution, it is clearly seen that the concentration of only either toxin or antitoxin RNA can be larger than zero.

After the plasmid loss, the dynamics of the active toxin mRNA and antitoxin sRNA are given in a simple form if we choose the concentrations at the steady state as the initial condition. They are divided into two cases shown below

**case I:** 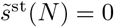

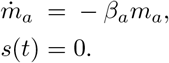
**case II:** 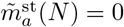

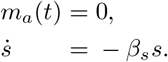

It is obvious that in the case II the concentration of the toxin protein stays zero after the plasmid loss. Thus, we study whether the concentration of toxin protein to have a peak value after the plasmid loss.

From (C.1), we get 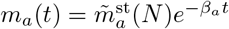, and the dynamics of the toxin protein is given by

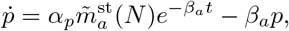

accordingly. The concentration of the toxin protein at time *t* is given as

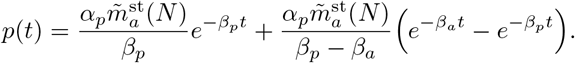

By differentiating *p*(*t*) respect to *t*, it is seen that *p*(*t*) is monotonically decreasing function regardless of parameter values, indicating that there is no peak of the toxin protein concentration during the plasmid loss.

## Appendix D. Analytical approximation of the peak concentration for a plasmid decrease event

For the quantitative estimate of the peak value of the toxin protein for each plasmid decrease event, we recall the timescale separation between the dynamics of the active toxin mRNA and antitoxin sRNA. This timescale separation allows us to assume that the concentration of the antitoxin sRNA quickly respond the change of the plasmid copy number (*N* → *N*’), and s relaxes to *s*^st^(*N*’) instantaneously. Because *ks*^st^(*N*’) ≫ *β_a_* holds, we ignore the sponaneous degradation term, and thus, here we use the approximated ones for the steady concentration of the molecules. In addition, the time-dependent solution of the inactive toxin mRNA is exactly calculated. Thus, we get an approximated differential equation for *m_a_* given by

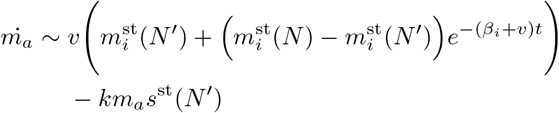

where 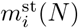 and *s*^st^(*N*’) represent the steady concentration under the given number of plasmid copies indicated in the parenthesis. By substituting the steady concentrations with the *β_a_* = 0 assumption (8) to this equation, we get

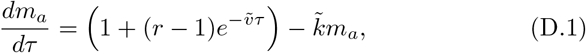

where 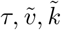, and *r* are defined as follows: 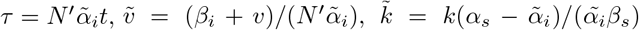, and *r* = *N/N*’. By solving (D.1), we get

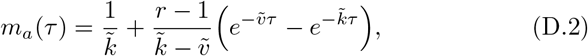

and the time *τ*\* at which *m_a_* reaches its peak is calculated as 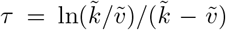. By substituting *τ*\* into (D.1) and setting its left-hand side to zero, the peak value of *m_a_* is obtained as

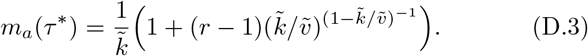

The ratio 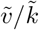 is given by a different form as 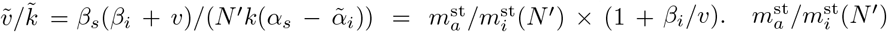 should be much less than one for the postsegrefational-killing mechanism to work, and *β_i_* and *υ* are in the same order of magnitude. Interestingly, *υ* ≈ *β_i_* gives the highest peak in varying *υ*. Thus, we set the ratio 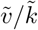 as zero (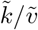 as infinity) leading to

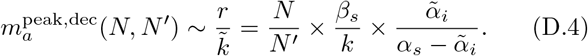

The maximum concentration of the toxin protein is also roughly estimated as 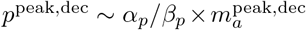 accordingly^+^.

As shown in figure D1, the peak value of the active toxin mRNA is approximately independent to the absolute number of the plasmid copies, whereas for large *N/N*′ region, the peak value is not determined only by the ratio *N/N*′ and deviates from the estimated value. This deviation happens because the assumption for the dynamics of the antitoxin sRNA is no longer valid for large *N/N’* region. We assumed that the concentration of the antitoxin sRNA quickly reaches to the new steady state with *N*′ plasmid copies (*s*^st^(*N*′)) at *t* = 0. For large *N/N*′ region, however, the concentration stays at values much lower than the steady state value, because the huge amount of the active toxin mRNA is still produced from the “reservoir” while the production rate of the antitoxin sRNA has dropped to the new production rate with *N*’ plasmid copies. Since there is less antitoxin sRNA than the steady state, the larger peak of the active toxin mRNA than the estimated value appears.

## Appendix E. The master equation for the model of type-I TA systems on bacterial chromosomes

We introduced the ON/OFF dynamics of the toxin gene transcription, and write down the model by using master equation to consider the stochasticity of chemical reactions.

*Q_L_* = *Q_L_*(*n_i_,n_a_,n_s_,n_p_;t*) represents the probability of state at which there are *n_i_* inactive toxin mRNA, *n_a_* active toxin mRNA, *n_s_* antitoxin sRNA, *n_p_* toxin proteins, and the toxin gene locus is in the Lth state (*L* = 0 and *L* =1 corresponds to the OFF and ON state, respectively) at the time *t*. Then, the master equation for the model is given as

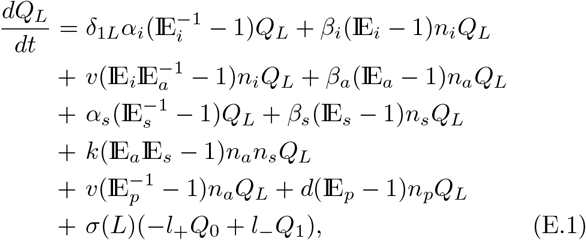

where 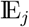 is a step operator of chemical species *j* which acts as 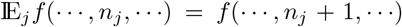. *δ_ab_* is Kronecker’s delta, and *σ*(*L*) is sign function of L which is given as *σ*(0) = 1 and *σ*(1) = −1.

For the model lacking the inactive toxin mRNA and the processing reaction, the probability of the states is defined as *Q_L_* = *Q_L_*(*n_a_, n_s_, n_p_;t*), and the master equation is slightly modifies as

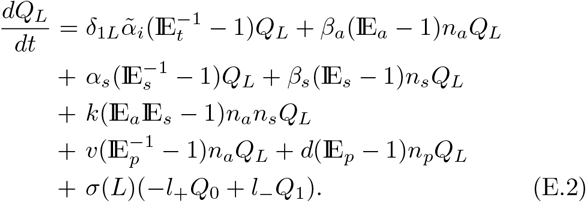

## Appendix F. Calculation of the noise intensity of translationally active toxin mRNA amount

Here, we show the models to obtain the Fano factor of the number of the active toxin mRNA. Since the pair degradation of the active toxin mRNA and the antitoxin sRNA and the translation of the active toxin mRNA are shared with the models with- and without the intermediate step (the inactive toxin mRNA and the slow processing reaction), it is not likely to have any roles in the difference in the noise in the number of the toxin protein. Thus, we removed the antitoxin sRNA and the toxin protein, and accordingly all the reactions which relate to these variables. This results in two simple models, namely the model consists of the inactive and active toxin mRNA, and the model consists of only the active toxin mRNA given as

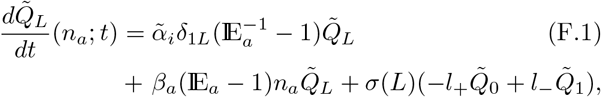

for the model without the intermediate step and

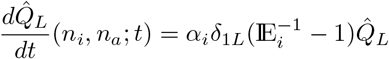

**Figure D1.**
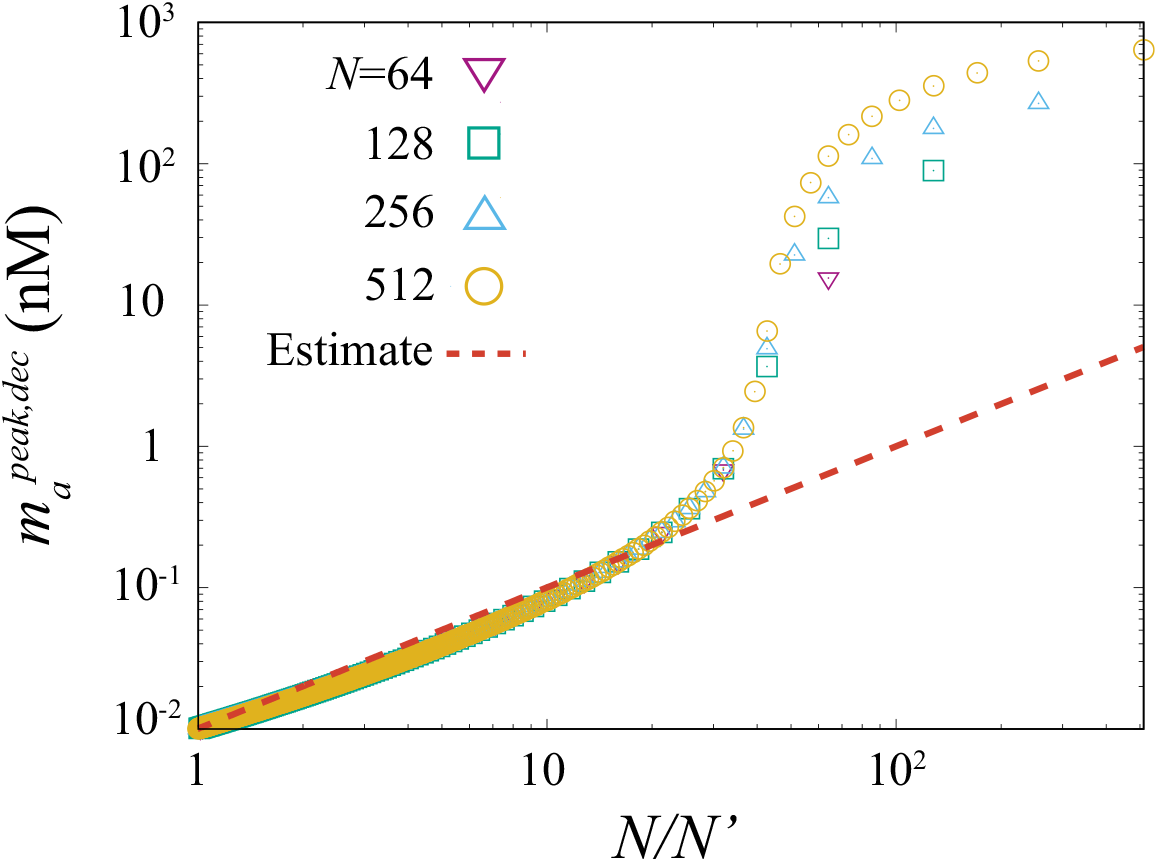
A comparison between the estimated value (D.3) and numerical results of the peak concentration of the active toxin mRNA for large *N* and *N*’. Numerical data are generated by the same method with figure 4 in main text.

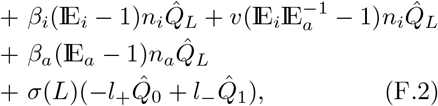

for the model with the intermediate step. 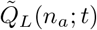 and 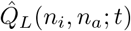 represent the probabilities of the indicated states.

Here, we introduce the generating functions defined as

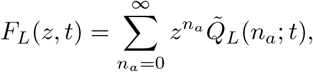

for (F.1), and

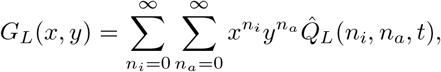

for (F.2).

We solved the equations for the generating functions in steady-state with the normalization condition 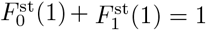 and 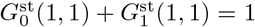. We obtained (12) in the main text with the coefficients given by

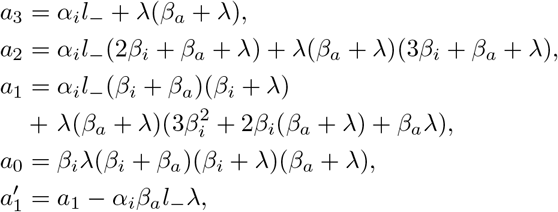

where λ is defined as λ = *l*_+_ + *1*_−_. It can be easily seen that all the five coefficients have positive values and 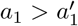 holds regardless of parameter values.

## Footnotes

‡ This rate is about 10-fold larger than the value obtained for *in vitro hok/sok* system [16]. The *in vivo* reaction rate is unknown for *hok/sok* system, but the values used in this paper are consistent with other *in vivo* estimates of sRNA-mRNA interactions [22].

§ This condition is typically satisfied if the parameters are chosen so that the PSK mechanism works. With such parameter values, *γ* ~ 1 (otherwise, the ratio between the peak- and steady concentration of the active toxin mRNA is small), hence the term *β_a_*(*γ* − 1)/(*ks*^st^(*N*)) is much smaller than 1.

∥ For larger *N/N*′ (*N/N*′ ≥ 40), the peak value also depends on the absolute number of the plasmid copies, see Appendix D

¶ The Fano factors obtained from (F.1) and (F.2) cannot become smaller than 1 regardless of parameter values, meaning that the inactive toxin mRNA and the slow processing reaction can reduce the noise at most to the same level as the simple Poisson processes.

+ The result (D.4) itself is also obtained by a simple calculation as following: First we assume that the dynamics of *m_i_* is much slower than that of *m_a_* so that 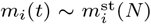 holds within a certain timescale, and approximate *dm_a_/dt* (D.1) as 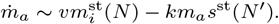 In addition, by regarding that the peak value of *m_a_* is obtained by setting 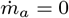, we get 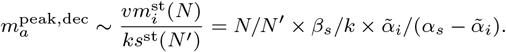

## Reference

[1] Pandey D and Gerdes K 2005 Nucleic Acids Research 33 966–976

[2] Unterholzner S J, Poppenberger B and Rozhon W 2013 Mobile genetic elements 3 e26219

[3] Page R and Peti W 2016 Nature chemical biology 12 208

[4] Harms A, Brodersen D E, Mitarai N and Gerdes K 2018 Molecular cell

[5] Gerdes K, Rasmussen P B and Molin S 1986 Proceedings of the National Academy of Sciences 83 3116–3120

[6] Verstraeten N, Knapen W J, Kint C I, Liebens V, Van den Bergh B, Dewachter L, Michiels J E, Fu Q, David C C, Fierro A C et al. 2015 Molecular cell 59 9–21

[7] Dörr T, Vulić M and Lewis K 2010 PLoS biology 8 e1000317

[8] Gerdes K, Helin K, Christensen O W and Løbner-Olesen A 1988 Journal of molecular biology 203 119–129

[9] Gong C C and Klumpp S 2017 PloS one 12 e0169703

[10] Gerdes K, Nielsen A, Thorsted P and Wagner E G H 1992 Journal of molecular biology 226 637–649

[11] Franch T, Gultyaev A P and Gerdes K 1997 Journal of molecular biology 273 38–51

[12] Gerdes K 2016 Phil. Trans. R. Soc. B 371 20160189

[13] Gultyaev A P, Franch T and Gerdes K 1997 Journal of molecular biology 273 26–37

[14] Gerdes K and Wagner E G H 2007 Current opinion in microbiology 10 117–124

[15] Arnion H, Korkut D N, Masachis Gelo S, Chabas S, Reignier J, Iost I and Darfeuille F 2017 Nucleic acids research 45 4782–4795

[16] Thisted T, Sørensen N, Wagner E and Gerdes K 1994 The EMBO journal 13 1960–1968

[17] Faridani O R, Nikravesh A, Pandey D P, Gerdes K and Good L 2006 Nucleic acids research 34 5915–5922

[18] Berghoff B A and Wagner E G H 2017 Current genetics 63 1011–1016

[19] Levine E, Zhang Z, Kuhlman T and Hwa T 2007 PLoS biology 5 e229

[20] Mitarai N, Andersson A M, Krishna S, Semsey S and Sneppen K 2007 Physical biology 4 164

[21] Mehta P, Goyal S and Wingreen N S 2008 Molecular systems biology 4 221

[22] Mitarai N, Benjamin J A M, Krishna S, Semsey S, Csiszovszki Z, Massé E and Sneppen K 2009 Proceedings of the National Academy of Sciences 106 10655–10659

[23] Sneppen K 2014 Models of Life (Cambridge University Press)

[24] Milo R and Phillips R 2015 Cell Biology by the Numbers (Taylor & Francis Group) ISBN 9781317230694

[25] Berghoff B A, Hoekzema M, Aulbach L and Wagner E G H 2017 Molecular microbiology 103 1020–1033

[26] Vogel J, Argaman L, Wagner E G H and Altuvia S 2004 Current Biology 14 2271–2276

[27] Giese K C, Michalowski C B and Little J W 2008 Journal of molecular biology 377 148–161

[28] Gillespie D T 1977 The journal of physical chemistry 81 2340–2361

[29] Gelens L, Hill L, Vandervelde A, Danckaert J and Loris R 2013 PLoS computational biology 9 e1003190

[30] Cataudella I, Sneppen K, Gerdes K and Mitarai N 2013 PLoS computational biology 9 e1003174

[31] Kawano M, Oshima T, Kasai H and Mori H 2002 Molecular microbiology 45 333–349

[32] Yamaguchi Y, Tokunaga N, Inouye M and Phadtare S 2014 Journal of molecular microbiology and biotechnology 24 91–97

